# Killer Immunoglobulin-like Receptor-Ligand Interactions Predict Clinical Outcomes Following Unrelated Donor Transplants

**DOI:** 10.1101/680298

**Authors:** Elizabeth Krieger, Roy Sabo, Sanauz Moezzi, Caitlin Cain, Catherine Roberts, Pamela Kimball, Alden Chesney, John McCarty, Armand Keating, Rizwan Romee, Christina Wiedl, Rehan Qayyum, Amir Toor

**Affiliations:** Massey Cancer Center, Virginia Commonwealth University; Department of Biostatistics, Virginia Commonwealth University; Department of Surgery, Virginia Commonwealth University; Department of Pathology, Virginia Commonwealth University; Princess Margaret Cancer Center, University of Toronto; Dana Farber Cancer Center, Harvard University; Department of Internal Medicine, Virginia Commonwealth University

## Abstract

Killer immunoglobulin-like receptor (KIR) and KIR-ligand (KIRL) interactions play an important role in natural killer (NK) cell mediated graft versus leukemia effect following hematopoietic cell transplantation (HCT). However, there is considerable heterogeneity in the KIR gene and KIRL content in individuals, making it difficult to estimate the full clinical impact of NK cell reconstitution following HCT. Herein a novel mathematical model designed to quantify these interactions is presented to better assess the influence of NK cell-mediated alloreactivity on transplant outcomes. Ninety-eight HLA matched unrelated donor (URD) HCT recipients were retrospectively studied. The KIR-KIRL interactions were quantified using a system of matrix equations. Unit values were ascribed to each KIR-KIRL interaction and directionality of interactions was denoted by, either a positive (activating) or negative symbol (inhibition); these interactions were then summed. The absolute values of both the missing KIRL as well as inhibitory KIR-KIRL interactions were significantly associated with overall survival and relapse. These score components were initially used to develop a weighted (w-KIR Score) and subsequently a simplified, non-weighted KIR-KIRL interaction scores (IM-KIR Score). Increased w-KIR Score and IM-KIR Score were both predictive of all-cause mortality and relapse; w-KIR score HR of 0.37 (P=0.001) and 0.44 (P=0.044) respectively; IM-KIR score HR of 0.5 (P=0.049) and 0.44 (P=0.002) respectively. IM-KIR score was also associated with NK cell reconstitution post HCT. KIR-KIRL interactions as reflected by the w-KIR and IM-KIR scores influence both relapse risk and survival in recipients of HLA matched URD HCT with hematological malignancies.

**Highlights:**

- Killer immunoglobulin-like receptor (KIR) – KIR-ligand (KIRL) interactions display a high degree of variability in HLA matched stem cell transplant (SCT) recipients.
- Individually, the magnitude of inhibitory and missing KIR-KIRL interactions predicts overall survival and relapse risk following SCT.
- Taken together the known KIR-KIRL interactions also predict clinical outcomes after HLA matched unrelated donor stem cell transplantation.

## Introduction

Hematopoietic cell transplantation (HCT) provides curative therapy for high-risk hematological malignancies^1^; however, transplant related mortality rates are high due to infection^2^, graft versus host disease (GVHD)^3^ and disease relapse^1^. Therapeutic benefit of a stem cell allograft is predominantly mediated through the alloreactivity of donor immune effectors directed at a recipient’s malignant cells and is termed the graft versus leukemia (GVL) effect^4^. Natural Killer (NK) cells are the first immune effector cells to reconstitute after allogeneic HCT, and are capable of affecting GVL ^5,6^; largely through germline-encoded receptors expressed on the NK cells, and inherited independently of human leukocyte antigens (HLA)^7^. These properties give NK cells a unique advantage, allowing them to mediate early GVL effects in a HLA matched environment, prior to the emergence of T cell-mediated GVL, without causing GVHD.

Human NK cells possess a multitude of different cell surface receptors classes; of these, the largest and most well studied are the killer immunoglobulin-like receptors (KIR). KIRs transduce either inhibitory or activating signals to the NK cell after interacting with, or in the absence of interactions with HLA and HLA-like ligands on the target cell surface^8^. The balance of these signals may lead to inhibition, or activation of the NK cell and target cell destruction through multiple mechanisms, including the release of cytotoxic granules containing mediators like perforin and granzyme^9^. Previous studies evaluating the role of KIR and HLA interactions in NK cell alloreactivity and HCT outcomes have included either one KIR-KIR ligand interaction at a time or have considered the donor KIR haplotypes. For example, missing KIR ligand interactions occur when the donor possesses inhibitory KIR (iKIR) for which the recipient lacks the corresponding HLA KIR ligand (KIRL)^10^. This missing KIRL (mKIRL) effect was shown to decrease relapse in haploidentical transplant recipients with acute myeloid leukemia (AML)^11^.

The KIR gene locus is highly polymorphic and has been classified into 2 haplotypes based on KIR gene content^7^, haplotype B containing a larger complement of activating KIR (aKIR) and haplotype A containing only one activating gene. HCT transplantation with KIR haplotype B donors are generally thought to yield favorable outcomes with relapse reduction, when compared to donors with KIR haplotype A, possibly due to the increased NK activation potential and greater GVL capabilities^9^. Along these lines, single activating KIRs have also been studied in relation to the recipient’s HLA status. It has been reported that relapse risk for AML is reduced when recipients with a HLA C1^+^ phenotype are transplanted using donors with activating KIR2DS1^12^. While these clinical associations are well characterized, as more evidence has been gathered, conflicting data have emerged^13–17^, in some instances disputing the NK cell mediated alloreactivity in HLA-matched HCT. Further, these studies have not fully accounted for the variability in donor KIR gene complement and recipient HLA types. It is very likely that this variability introduces a high degree of heterogeneity in donor NK cell-recipient target cell interactions. The lack of knowledge regarding these interactions compromises optimal donor selection for allogeneic HCT. This lack of clarity may stem from the analytic approach employed in conducting these studies, which has hitherto fore been similar to that taken in most transplant outcomes research, that is, correlation of specific biological characteristics (KIR genotype) with clinical outcomes. Herein, a novel analytic method which quantifies the interactions which might occur between variables controlling NK cell function is proposed to develop a system of scores which mathematically quantify the cumulative KIR-KIRL interactions in individual transplant recipients. In this system, missing KIR ligand, and the inhibitory and activating KIR-KIR ligand interactions are considered summative in their effect in mediating NK cell influence on clinical outcomes. Such a scoring system may allow better prediction of NK cell mediated GVL effect that may be expected from different HLA-matched donors for the same recipient. These scores -- if validated in a large cohort of patients -- may be used to select optimal HCT donors that yield an adequate GVL effect.

## Methods

### Patients

Virginia Commonwealth University (VCU) Institutional Review Board gave approval to conduct this retrospective study. All 8/8 HLA-matched unrelated donor-recipient pairs (DRP) who had KIR genotyping performed and were transplanted at VCU between 2014 and 2017 were retrospectively studied. Absolute NK cell counts were measured on days +30, +60 and +100 post-transplant by flow cytometry of peripheral blood samples utilizing antibodies against CD56 and CD3 antigens and the NK cells were defined as CD3−/CD56+. Acute and chronic GVHD were assigned utilizing the Glucksberg and NIH consensus criteria, respectively.

### KIR and KIR Ligand Assignment

Patients and their donors were matched at 8/8 loci, including HLA-A, HLA-B, HLA-C, HLA-DRB1. High throughput HLA sequence-based typing was performed after DNA was isolated from blood or buccal samples (Protrans, Ketschau, Germany). KIR genotyping was determined by intermediate resolution, quantitative polymerase chain reaction (Immucor, Norcross, GA) using LinkSeq KIR 384 (One Lambda Canoga Park, CA). KIR genotyping was not considered during donor selection. HLA epitopes for HLA-B and HLA-C recognized as KIRL by KIR were determined using the European bioinformatics KIR Ligand calculator (https://www.ebi.ac.uk/ipd/kir/ligand.html). Every HLA-C allotype was designated as either C1 or C2. Similarly, HLA-B allotypes were also divided into 2 epitopes Bw4 and Bw6. Bw4 is a KIR epitope, as are HLA-A3 and -A11. KIR and KIRL interactions were determined as described in supplementary table 1 (adapted from S. Cooley et al 2018^9^).

### KIR-KIRL interaction scores

To mathematically derive the KIR-KIRL interaction score, the interactions were viewed from the frame of reference of the donor NK cells (Figure 1). KIR-KIRL interaction values were assigned in the following manner: if an inhibitory KIR (iKIR) had a ligand, this resulted in an interaction (−1) × (1) = −1, which gave the NK cell an inhibitory signal. If the inhibitory KIR did not have its ligand (mKIRL), its inhibitory effect is abrogated, and since it is assumed that under basal conditions NK cell are constitutively active, this was be described by the interaction (−1) × (−1) = +1. Interactions between activating KIR (aKIR) and their ligands analogously were given, (1) × (1) = +1 when the ligand was present and (1) × (0) = 0, when the ligand was absent. In the latter instance, the absence of a ligand is characterized by a 0 rather than −1, because rather than an inhibitory signal being abrogated by the absence of its ligand, in this instance the activating signal is simply not given since the aKIRL is not present (Figure 1). With these values assigned to each KIR-KIRL interaction, the aggregate KIR effect on the NK cells may be quantified by a system of matrix, vector-operator equations (adapted from Abdul Razzaq et al, 2016^18^ and Koparde et al, 2017^19^). In this system the multicomponent, NK cell-KIR-vector is composed of the KIRs which recognize specific KIRL present on the target cell-operator, which transforms the afore-mentioned vector. The total magnitude of these interactions may be derived by a matrix multiplication operation (absence of an aKIR cognate ligand is designated noKIRL in this system),

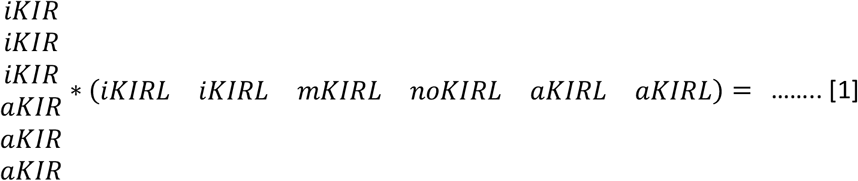

Solving this equation yields

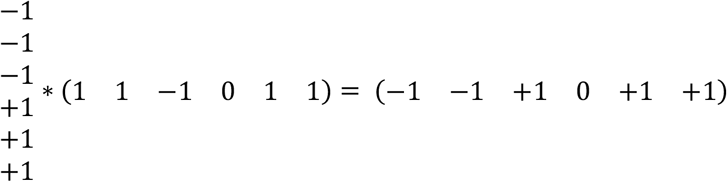

**Figure 1.**
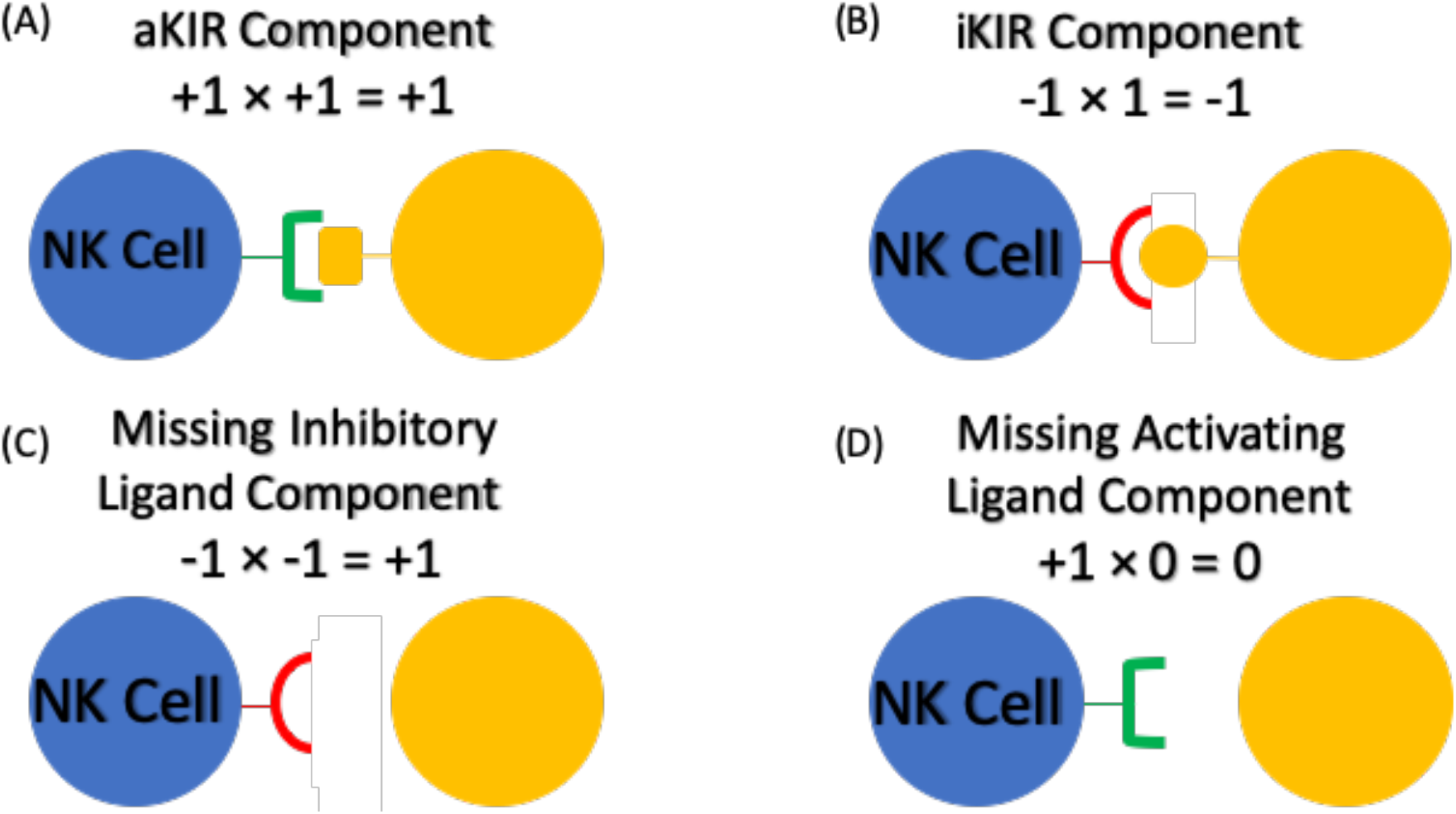
NK cell - Target Cell interactions were assigned values (A) Donor cell with activating KIR ligand (aKIR) and recipient cell with corresponding HLA ligand (B) Donor cell with Inhibitory KIR ligand (iKIR) and recipient cell with corresponding HLA ligand (C) Donor cell with iKIR and recipient cell without corresponding HLA ligand (missing ligand) (D) Donor cell with aKIR and recipient cell without corresponding HLA ligand

The total KIR-KIRL interaction score in this model is the sum of all these interactions, and in this instance equals 1. These scores were calculated for all donor-recipient pairs. Next, these scores were resolved into their components, as in vector addition interactions (Supplementary Figure 1). In other words, while the total magnitude and direction of the KIR-KIRL interaction represents a sum of the activating, inhibitory and missing KIR-KIRL interactions, these components may also be considered individually. These components of the total KIR-KIRL interaction scores will then give an estimate of the NK cell mediated alloreactivity resulting from a specific class of interactions. To accomplish this, the total inhibitory KIR score was calculated, and the absolute value (designated as **|**…**|**) of the interaction between iKIR (vector) and the corresponding KIRL in the (operator) was determined

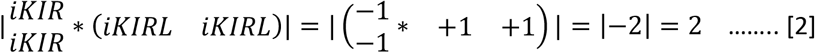

Total aKIR score component was similarly calculated by taking the absolute value of the interaction between aKIR (vector) and corresponding KIRL in the (operator)

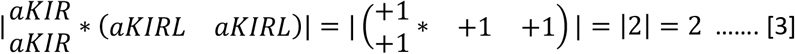

The missing KIRL component was computed by taking the absolute value of the product of the iKIR (vector) present without the corresponding KIRL in the (operator)

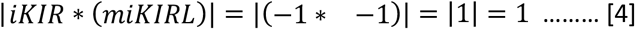

### Statistical Methods

Time to relapse and death were determined from the day of transplant. Associations between KIR-KIRL interaction scores or the score components outlined above, and time-to-event outcomes (relapse and mortality) were estimated using parametric survival analysis. Given the exploratory nature of the work with this novel scoring system, the choice of distribution which minimized the Bayesian Information Criterion (or BIC) was chosen from either exponential, Weibull, or gamma distributions; models were fit separately for each outcome and for the KIR score or its components. KIR-KIRL component score models were also fit in both unadjusted and adjusted manners, where in the latter case the models included CD34+ cell dose infused, recipient age at transplantation, recipient sex, conditioning intensity, whether ATG was administered or not, donor KIR haplotype, myeloid disease, and disease status at transplant. The UNIVARIATE, FREQ, CORR and GLIMMIX procedures from the SAS statistical software (version 9.4, Cary, NC, USA) are used for all KIR-KIRL component summaries and analyses.

Next, unique weights for inhibitory and activating KIR, and missing KIR ligand score components were generated separately to determine relapse-free survival using Cox proportional hazards models. These weights were used to generate weighted donor-recipient KIR-KIRL interaction scores for each individual to predict all-cause mortality and relapse. Subsequently, the two clinical endpoints (mortality and relapse) were studied by generating a single score by the unweighted addition of total inhibitory KIR and total missing ligand scores. Association of weighted and unweighted scores with both mortality and relapse, were examined using Cox proportional hazards models. To further examine the significance of these weighted and unweighted scores, we examined their association with NK cell count using mixed linear models with unstructured covariance. These analyses were performed using Stata 14.1 for MS Windows (StataCorp. 2015: Release 14. College Station, TX: StataCorp LP).

## Results

### Demographics

The study cohort comprised 98 patients who underwent 8/8 HLA matched unrelated donor HCT for hematologic malignancy (Table 1). KIR gene frequencies within our population were as follows: 97% KIR-2DL1, 49% KIR-2DL2, 93% KIR-2DL3, 94% KIR-3DL1 ^20^. KIRL frequencies were: 70% for C1, 84% for C2, 40% for Bw4, and 22% for both A3 and A11.

**Table 1.**
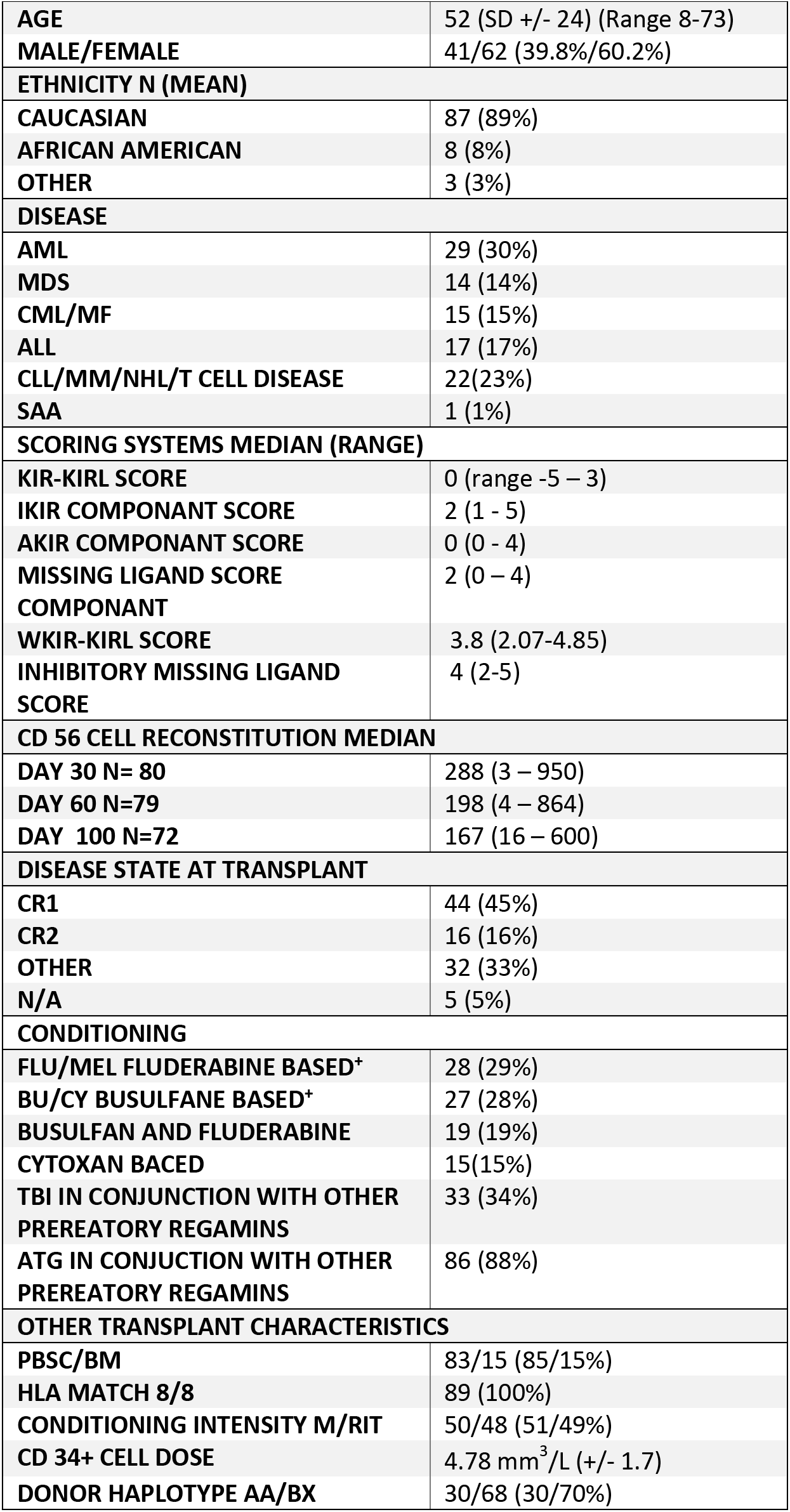
Patient characteristics, details of conditioning regimen, and CD56 reconstitution (N=98).

### KIR-KIRL interaction score

Total KIR-KIRL interaction scores (Equation 1) ranged between −5 to +3, with a median of 0 (Figure 2) and a distribution approximating a normal distribution. There was no significant difference in the total KIR-KIRL interaction scores between donors with KIR haplotype A/A or B/x. This variability in the derived KIR-KIRL interaction scores implies that within HLA identical donors there is considerable heterogeneity in the potential for NK cell mediated alloreactivity. Relationships between NK cell reconstitution and this score were explored in a subset of DRPs who had NK cell counts measured at days 30, 60, and 100 post-transplant. A linear relationship between the total KIR-KIRL score and NK cell count recovery was demonstrated in the patients examined (Figure 3A). When divided into 3 groups based on the magnitude of the score -- i.e., with negative (scores ranging from −5 to −3), a neutral (−2 to 0) or positive scoring group (1-3) -- the NK cell counts were significantly different between the three score groups. When compared to the lowest scoring group which had the highest NK cell counts, the neutral scoring group (p=0.049) and high scoring groups (p=0.013) had significantly lower counts (Figure 3B). These data suggest a relationship between the magnitude of KIR-KIRL interaction post-transplant and NK cell reconstitution.

**Figure 2.**
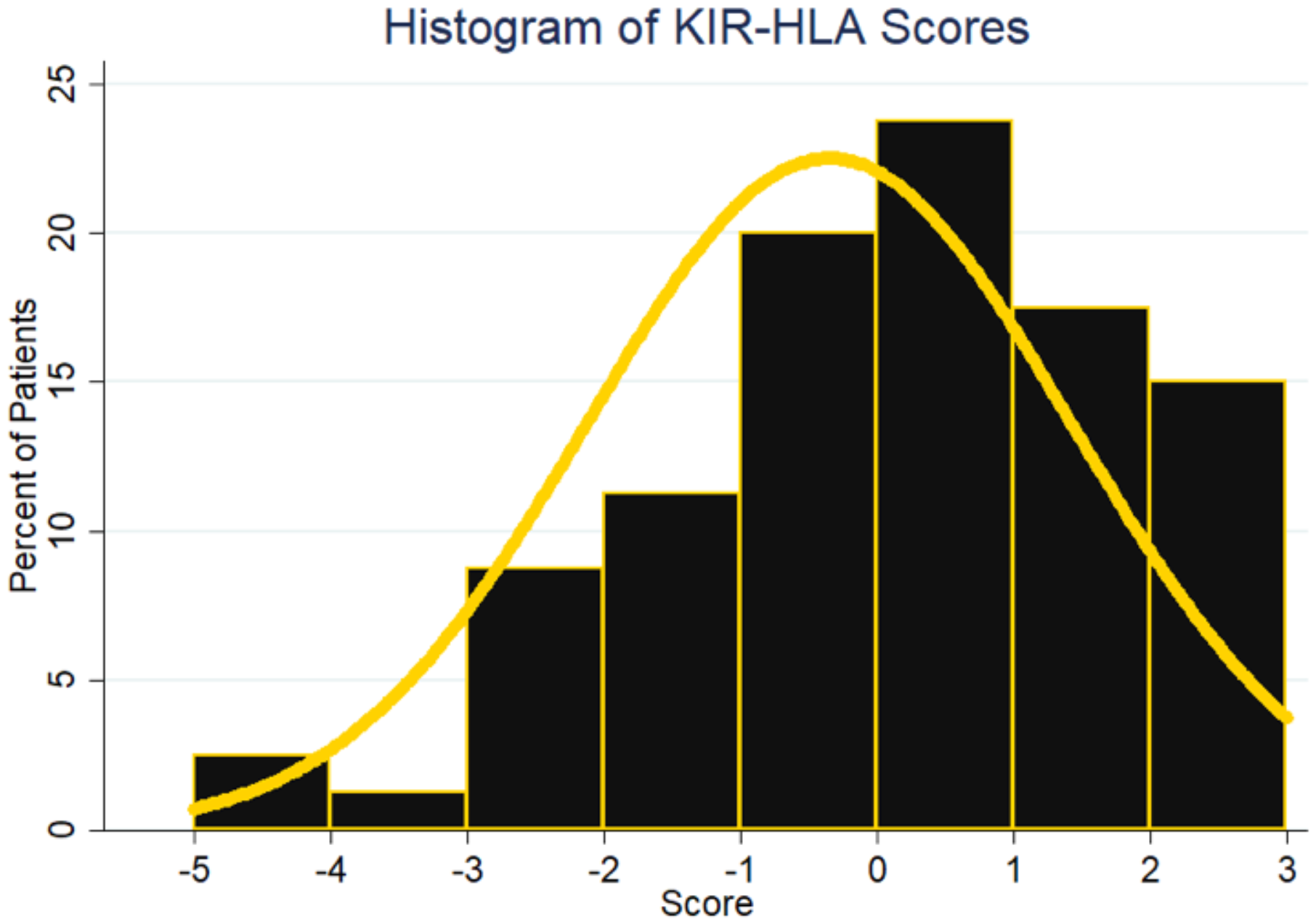
Frequency histogram utilized to represent the distribution scores for the 98 patient cohort when scored using the KIR-KIRL interaction scoring system. Yellow line used to represent an overlay of normal distribution

**Figure 3.**
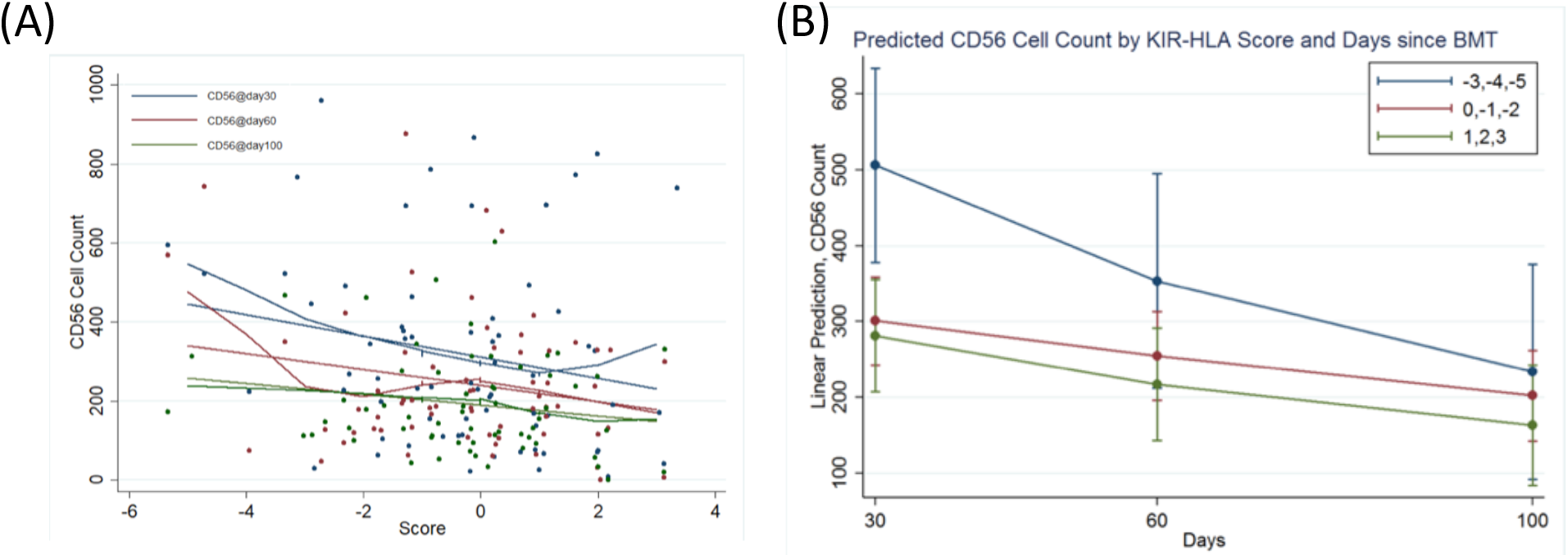
CD56 + cell count kinetics and KIR-KIRL interaction score (A) represents a scatter plot of CD56 + cells at day 30 (blue) day 60 (red) and day 100 (green) with a local regression, LOWESS line representing CD56 + cell counts with KIR-KIRL interaction score. (B) Predicted CD56 cell counts by KIR-KIRL interaction score and days since HSCT. the lowest KIR-KIRL interaction scorers (−5 to −3) had the highest CD56 reconstitution at day 30.

### KIR-KIRL interaction score components and survival

Given the relationship between KIR-KIRL scores and NK cell reconstitution any association between these scores and clinical outcomes was explored. While the total KIR-KIRL score did not have a significant impact on overall survival, there was a trend suggesting that components of the score might influence outcomes. KIR-KIRL score components were then examined for their impact on all-cause mortality. Both the inhibitory KIR-KIRL interaction score (Equation 2) and the missing KIRL score (Equation 4) were significantly associated with protection from all-cause mortality, with a hazard ratio of 2.7 (95% CI: 1.5, 4.6; p-value = 0.0006) and 3.2 (95% CI: 1.7, 6.0; p-value = 0.0004) respectively. However, the activating KIR component score (Equation 3) did not have this protective effect (p-value = 0.4126). These results demonstrated a positive association between the score component magnitudes and time to death, i.e., the larger the magnitude, the longer patients survived. When adjusting for other measures, (Table 2) both the inhibitory KIR-KIRL interaction score (HR = 2.1, 95% CI: 1.2, 3.6; p-value = 0.006) and the missing KIRL score (HR = 2.7, 95% CI: 1.5, 5.1; p-value = 0.001) remained associated with mortality.

**Table 2.** KIR-KIRL interaction score components unadjusted and adjusted hazard ratios for time to death.

| Table 2: Adjusted Hazard Ratios for Time to Mortality |  |  |  |  |  |  |  |
| --- | --- | --- | --- | --- | --- | --- | --- |
|  |  | Unadjusted |  |  | Adjusted |  |  |
| Measure |  | HR | 95% CI | p-value | HR | 95% CI | p-value |
| Inhibitory KIR |  | 2.7 | 1.5, 4.6 | 0.0006 | 2.1 | 1.2, 3.6 | 0.0059 |
| Activating KIR |  | 1.2 | 0.7, 2.0 | 0.4126 | 1.3 | 0.7, 2.4 | 0.3437 |
| Missing KIR |  | 3.2 | 1.7, 6.0 | 0.0004 | 2.7 | 1.5, 5.1 | 0.0015 |
| CD34 Dose |  |  |  |  | 1.3 | 1.0, 1.6 | 0.0319 |
| Age |  |  |  |  | 0.9 | 0.9, 1.0 | 0.0003 |
| Sex | F vs M |  |  |  | 1.1 | 0.5, 2.3 | 0.7909 |
| Treatment Intensity | M vs RIT |  |  |  | 1.4 | 0.7, 3.0 | 0.3214 |
| ATG | Y vs N |  |  |  | 0.8 | 0.2, 3.0 | 0.7373 |
| Donor Haplotype | A vs B |  |  |  | 1.6 | 0.7, 3.8 | 0.2928 |
| Myeloid Disease | Y vs N |  |  |  | 1.5 | 0.7, 3.2 | 0.2938 |
| Disease State | CR1 vs Other |  |  |  | 1.1 | 0.5, 2.4 | 0.8853 |

### KIR-KIRL interaction score components and risk of relapse and GVHD

KIR-KIRL score components were next studied for association with relapse. Both the inhibitory KIR-KIRL interaction score (Equation 2) and missing KIRL scores (Equation 4) were significantly associated with relapse prevention, demonstrating HR of 2.6 (95% CI: 1.3, 5.4; p-value = 0.01) and 3.5 (95% CI: 1.5, 8.3; p-value = 0.005) respectively. Activating score component (Equation 3) did not have a similar impact (p-value = 0.84). As noted above, these findings imply that the larger the KIR-KIRL interaction component score magnitudes, the less likely relapse is to occur and the longer it takes for it to occur. When adjusting for other measures (Table 3) the association for the inhibitory score was no longer significant (p-value = 0.37), though the missing KIR ligand score component remained significantly associated with relapse with a HR of 2.4 (95% CI: 1.0, 5.8; p-value = 0.04), again with a positive association with time to relapse. It is to be noted that the KIR-KIRL and KIR-KIRL component scores were not significantly associated with acute or chronic GVHD, or CMV reactivation. These data suggest that a higher inhibitory KIR and missing KIRL components conferred greater protection from relapse in HLA matched URD HCT.

**Table 3.** KIR-KIRL interaction score components unadjusted and adjusted hazard ratios for time to relapse.

| Table 3: Adjusted Hazard Ratios for Time to Relapse |  |  |  |  |  |  |  |
| --- | --- | --- | --- | --- | --- | --- | --- |
|  |  | Unadjusted |  |  | Adjusted |  |  |
| Measure |  | HR | 95% CI | p-value | HR | 95% CI | p-value |
| Inhibitory KIR |  | 2.6 | 1.3, 5.34 | 0.0106 | 1.4 | 0.7, 3.1 | 0.3754 |
| Activating KIR |  | 0.9 | 0.5, 1.8 | 0.8513 | 1.1 | 0.5, 2.8 | 0.7806 |
| Missing KIR |  | 3.5 | 1.5, 8.3 | 0.0047 | 2.4 | 1.0, 5.8 | 0.0443 |
| CD34 Dose |  |  |  |  | 1.6 | 1.1, 2.3 | 0.0106 |
| Age |  |  |  |  | 1.0 | 0.9, 1.0 | 0.3148 |
| Sex | F vs M |  |  |  | 1.0 | 0.3, 3.5 | 0.9458 |
| Treatment Intensity | M vs RIT |  |  |  | 6.3 | 1.8, 22.0 | 0.0043 |
| ATG | Y vs N |  |  |  | 1.6 | 0.3, 8.2 | 0.5715 |
| Donor Haplotype | A vs B |  |  |  | 3.2 | 0.6, 16.7 | 0.1589 |
| Myeloid Disease | Y vs N |  |  |  | 4.6 | 1.1, 20.0 | 0.0422 |
| Disease State | CR1 vs Other |  |  |  | 2.9 | 0.7, 12.8 | 0.1503 |

### Weighted KIR-KIRL interaction scores

Given the observed impact of the KIR-KIRL components on survival and relapse the relative weights of these interactions on mortality and relapse were calculated and a weighted total KIR-KIRL interaction score determined. The Cox model for mortality associations (Table 4) was utilized to derive an equation for a weighted KIR score (wKIR score) for each donor recipient pair

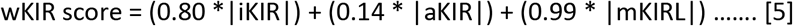

**Table 4.** KIR-KIRL interaction score components Cox Model Mortality associations.

| Table 4. Cox Model Mortality Associations |  |  |  |  |
| --- | --- | --- | --- | --- |
|  | Coefficient | Std. Err. | 95% Conf. Int | P Value |
| **iKIR-KIRL Component Score** | -0.80 | 0.27 | -1.32, -0.27 | 0.003 |
| **aKIR-KIRL Component Score** | -0.14 | 0.25 | -0.62, 0.35 | 0.58 |
| **mKIR-KIRL Component Score** | -0.99 | 0.31 | -1.6, -0.37 | 0.002 |

This equation demonstrates the differential impact of the various score components on the final score, the distribution of which was relatively broad in this cohort of 98 HLA matched unrelated DRP (Figure 4). The weighed KIR-KIRL interaction score again demonstrates considerable variability across HLA matched donor-recipient pairs, approximating a normal distribution. Reflecting the earlier findings with KIR component scores, this combined, weighted score (Equation 5) was predictive of all-cause mortality with a HR of 0.37 (95% CI: 0.2, 0.7, P=0.001). This implies that each unit increase in the weighted score results in a 63% decrease in the risk of all-cause mortality following HCT. This weighted score was also predictive of relapse risk with a HR of 0.44 (95% CI: 0.2, 1.0, P=0.044), indicating that for each unit increase in weighted total KIR-KIRL interaction score there was a 56% decrease in risk of relapse. These associations of weighted scores demonstrate the relative influence of the various components of the KIR-KIRL interaction scores with mortality and relapse risk following HCT. As in the previous component analysis, the inhibitory KIR and missing KIRL score components have the largest impact on relapse prevention.

**Figure 4.**
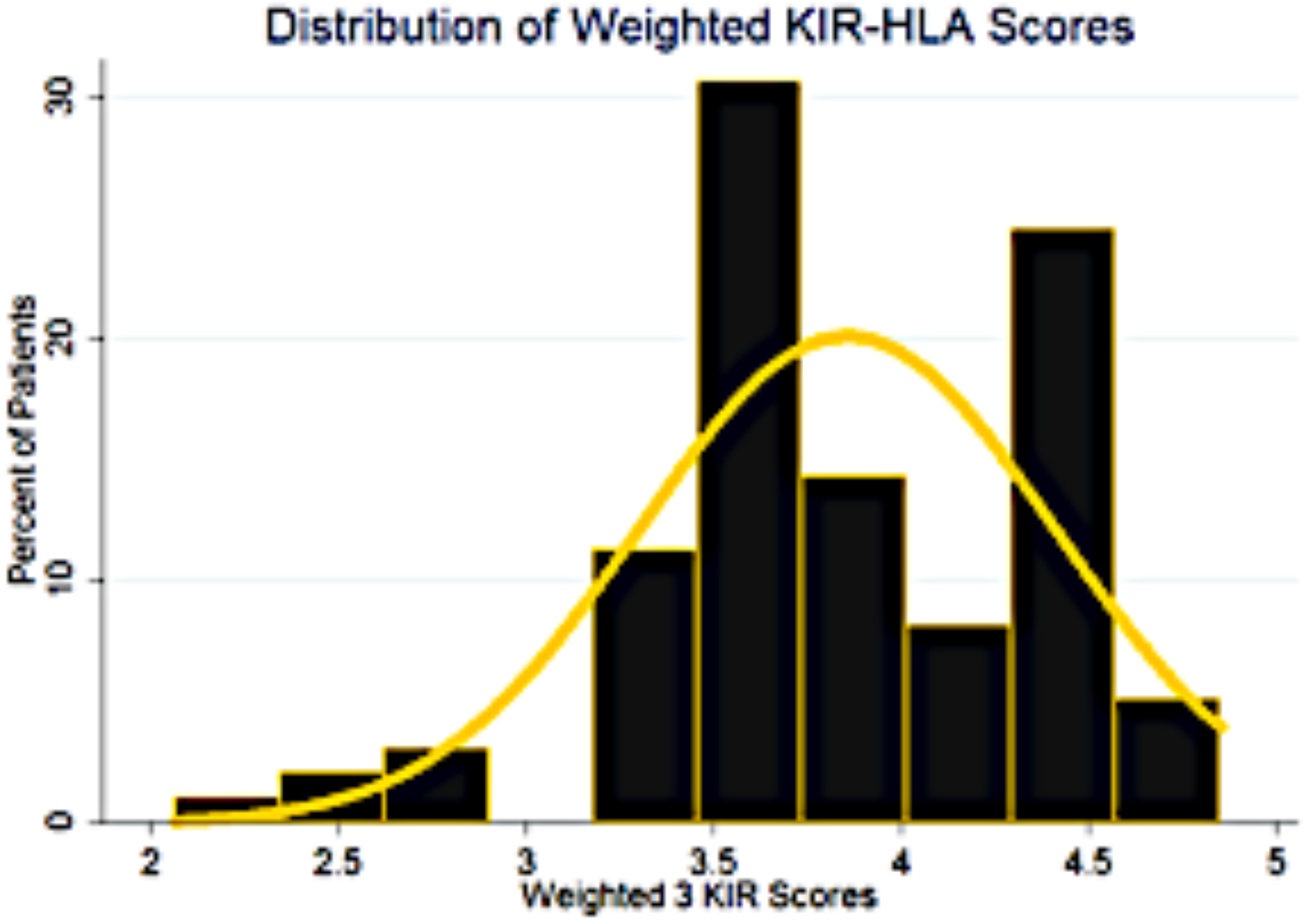
Frequency histogram utilized to represent the distribution scores for the 98 patient cohort when scored using the w-KIR scoring system. Yellow line used to represent an overlay of normal distribution

### A simplified 2 component score for predicting HCT outcomes

Given the influence of inhibitory KIR and missing KIRL score components (Equations 2 & 4; Table 4), a simple, non-weighted new score was developed using these heavily weighted score components. For these calculations both the components were given equal weights (Equation 6) to calculate the inhibitory-missing KIR ligand score (IM KIR Score)

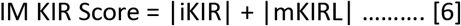

This IM KIR score took on the values 4.3±0.6 (Figure 5). Using Cox proportional hazards model, it was found that for each unit increase in the IM-KIR score, there was a 56% decrease in risk of all-cause mortality (HR = 0.44; 95%CI: 0.26 to 0.73; P=0.002), and a 50% decrease in risk of relapse (HR = 0.5; 95%CI: 0.25 to 1.0; P=0.049), confirming the importance of inhibitory KIR-KIRL and missing KIRL interactions in determining clinical outcomes following HCT, in the cohort reported here.

**Figure 5.**
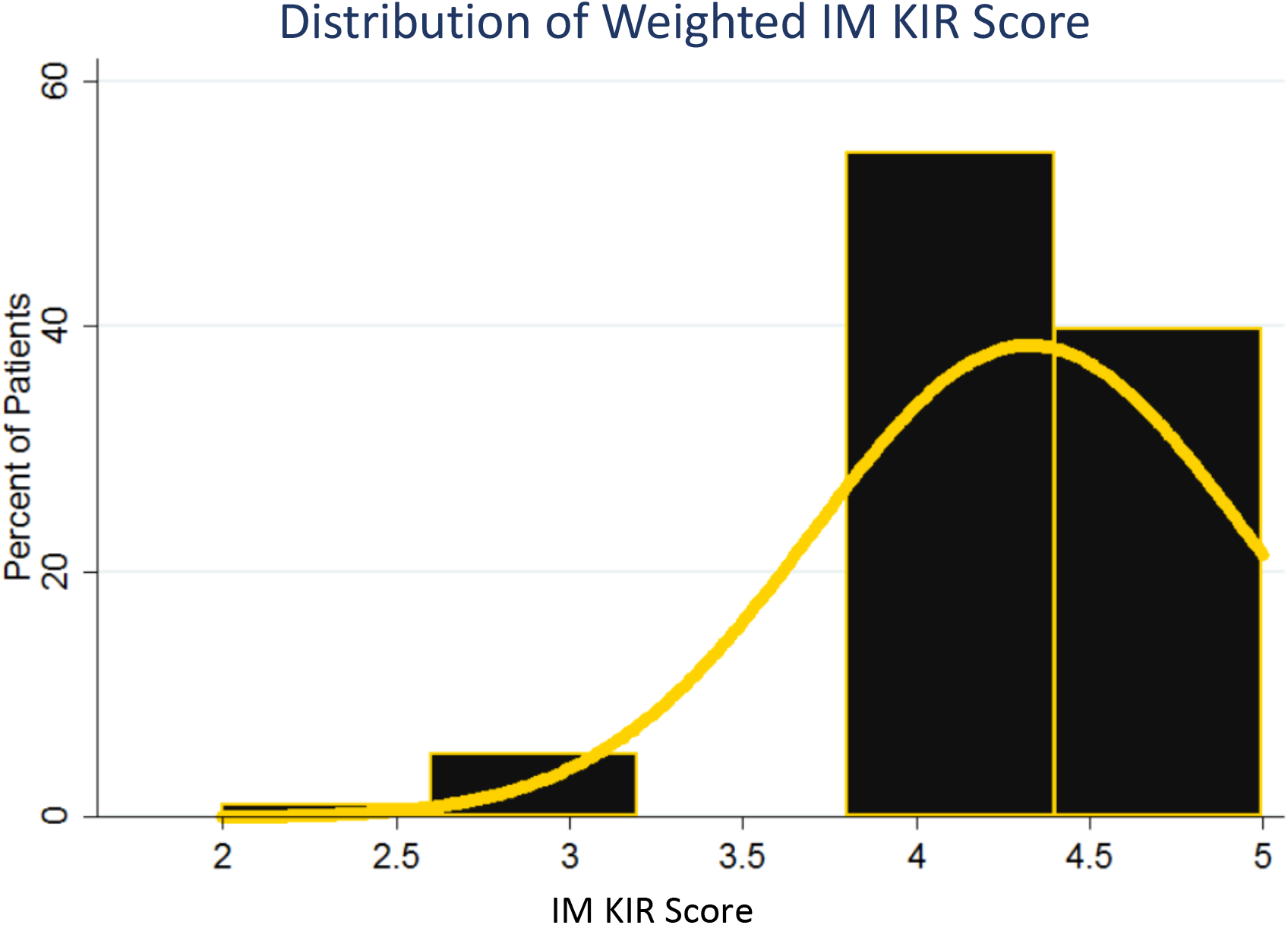
frequency histogram utilized to represent the distribution scores for the 98 patient cohort when scored using the IM KIR scoring system. Yellow line used to represent a normal distribution

### IM KIR Score’s association with NK cell count recovery

NK cell reconstitution was examined as a function of the IM KIR Score and demonstrated that an increase in the score of 1 unit was associated with an increase of fifty NK cells/μL (SE 25.2, P= 0.046). This suggests that KIR-KIRL interactions influence NK cell reconstitution after HCT.

## Discussion

This study was designed to develop a logic-based quantitative method to understand the variation in, and the cumulative effect of KIR and KIRL interactions on clinical outcomes following HLA-matched unrelated HCT. In doing so, this method departs from the conventional analytic methodology employed in examining the impact of pretransplant variables on clinical outcomes, where such characteristics are used to statistically derive probabilities of specific clinical outcomes, without consideration of direct interactions between these variables. The findings reported in this paper demonstrate that within a cohort of HLA identical patients with both myeloid and lymphoid malignancies, the magnitude of the KIR-KIRL interactions vary considerably, approximating a normal distribution. Further, both the inhibitory KIR-KIRL interactions, as well as the missing KIR ligands when mathematically determined are associated with mortality and relapse risk, albeit in a heterogeneous cohort of patients. When examined cumulatively in a patient cohort which primarily received ATG for GVHD prophylaxis, both a weighted total KIR-KIRL interaction score, as well as a non-weighted IM-KIR score (combining inhibitory KIR and missing KIRL interaction magnitudes) were similarly associated with improved survival and decreased relapse. These KIR-KIRL interactions are also associated with the magnitude of NK cell reconstitution. This novel formalized mathematical framework for quantifying KIR-KIRL interactions presented here may therefore be predictive of clinical outcomes in recipients of HLA matched unrelated donor allografts and may help identify optimal donors from amongst equally well HLA matched donors and merits further study in a larger cohort of patients.

The KIR gene locus shows a very high degree of variability, similar to the major histocompatibility locus encoding HLA molecules ^21^. In order to understand and quantify the effect KIR-KIRL interactions have on clinical outcomes following HCT, one must consider a model that accounts for variability in both these loci in the donors as well as the recipients. Genetic drift has provided a favorable distribution of KIR and HLA across populations as these molecules confer protection against pathogens encountered, facilitate favorable placental implantation^22^ and prevent autoimmune disease^20^. For example, the inhibitory KIR2DL2 gene is present in 46% of Europeans and 80% of Ethiopians while these populations are at least heterozygous for its ligand, C1 in 82% and 72% of the members, respectively. These patterns vary across the globe, such as in the Japanese population a far greater percentage of C1 KIRL with carrier frequencies of 96% and only 18% for C2^20^ is observed, with corresponding KIR2DL2 and KIR2DL3 frequency of 9% and 100% respectively. Conversely, HLA-B allotypes with epitope Bw4 and HLA-A3/A11 have much lower carrier frequencies world-wide, at 35% and 16% respectively^23^. However, their corresponding KIRs are far more common within European populations, which express carrier frequencies of 93% for the Bw4 epitope KIR3DL3 and 47% for the HLA-A11 epitope KIR2DS2^20^. This variation between KIR and KIRL population frequencies means that within HLA matched donor-recipient pairs there may be significant heterogeneity in NK cell mediated alloreactivity, and thus disease relapse and mortality risk. This was borne out in the analysis reported here where HLA matched donor recipient pairs demonstrated an approximately normal frequency distribution curve for the various KIR-KIRL interaction scores. Therefore, quantifying the magnitude of KIR-KIRL interaction reported here, may optimize the HCT donor selection process beyond HLA matching, particularly when it comes to NK cell mediated alloreactivity. It is to be noted that for this cohort KIR typing information was not utilized in donor identification, eliminating a source of bias.

The initial studies of NK cell alloreactivity using missing ligand analysis did not consider KIR gene variability. There are more than 30 KIR genotypes known^7^, which can be functionally split into 2 haplotypes, A and B as previously described^9^. This simple distinction was first used to look at the effect of additional aKIR gene content on clinical outcomes in HCT. The presence of KIR haplotype B alone was associated with reduced risk of relapse and increasing survival (RR relapse or death, 0.70 95% CI 0.55-0.88) in AML, however it did not have the same effect in ALL, and was not always reproducible^24^. This clinical benefit was then found to increase when donors homozygous for centromeric, B cen-specific gene content were utilized (RR relapse or death 0.85 95% CI 0.73-0.99). One theory for this phenomenon is the presence of the stronger binding affinity, KIR2DL2 and the absence of the low binding affinity, KIR2DL3. However others have reported reduced overall survival with higher B cen content^13^. And while it has been shown that allografts from donors positive for aKIR KIR2DS1 decrease the risk of relapse with a hazard ratio of 0.76, further studies on in vivo T cell depleted patients have shown an increased risk of relapse with increasing aKIR genes with a HR of 1.37. This model demonstrating increased variability in KIR-KIRL interactions may help understand some of these inconsistencies reported in the literature. In addition, the magnitude of effect reported here in a combined myeloid and lymphoid disease population far exceeds the effect previously reported in the KIR haplotype studies. One reason for this may be related to the loss of information that occurs when the haplotype is considered, as the expected HLA interactions are not accounted for, diminishing the signal strength one may get from KIR haplotype analyses. A mathematical frame work accounting for all the individual KIR-KIRL interactions is not susceptible to such loss of information.

The model reported here is robust in that its simple mathematical basis will allow incorporation of other NK cell receptors, alleles and their interactions, as additional data become available in the future. For example, after HCT CD56^bright^ NK cells are present at a high frequency, normalizing over time in the first year after transplant. This subset of NK cells express a lower level of KIR and a higher level of the inhibitory NK cell marker NKG2A:CD94^25^. NKG2A:CD94 is known to contribute to education of these early cells^26^. CD56^bright^ and CD56^dim^ percentages as well as NKG2A were not available for our cohort. However, moving forward this and other possible yet undiscovered NK cell-target cell interactions can be easily accounted for in the equations reported here.

This initial retrospective study was performed on a small cohort of patients with with known KIR typing available at a single institution. This cohort of is heterogeneous with multiple diagnoses, different conditioning regimens used and ATG used for GVHD prophylaxis in a majority of the patients. T cell depletion has been shown to alter immune cell reconstitution including, NK cell recovery dynamics^27,28^. Meaning that NK cell effect size may be larger in patients who receive *in vivo* or *ex vivo* T cell depleted grafts. Therefore, it is imperative that this analysis be repeated in a larger more homogeneous validation cohort, where adequate representation of both myeloid and lymphoid malignancy is ensured to have confidence that the findings reported here may be reliably reproduced. Data requests to accomplish this has been initiated through the CIBMTR data request process.

There are interesting questions raised by this analysis, one of which is, while KIR-KIRL score components are predictive of clinical outcomes, why is the unweighted total score in and of itself not so? When considering this question in mathematical terms, it is known that NK cells express their KIR in a stochastic fashion^21,29^, where within an individual’s NK cell repertoire some NK cells express no KIR, while others express every KIR within their genotype, as well as every combination of KIR expression in-between. This implies that there may be many NK cell ‘clones’ in each individual, each expressing a different KIR complement (see Supplementary material for mathematical supplement). The model reported herein scores the KIR-KIRL interaction score on the extreme end of this spectrum, assuming that all of the known KIR are expressed on all the NK cells which may dampen the effect of the individual score components if they are variably expressed between different NK cell clones.^30^ This implies that based on their expression, individual KIR may have to be weighted differently in the model reported here to improve its predictive value. In this analysis relative weights of different classes of KIR interactions determined by clinical correlation increased the predictive capacity of the total KIR-KIRL interaction scores, when the wKIR scores were evaluated.

The other question raised by these findings is why do inhibitory KIR have such a profound effect on clinical outcomes? To answer this in quantitative terms, one may take a dynamical systems view of NK cell responses as was previously done for T cell alloreactivity ^18,19^. In such systems, the future states of the system are dependent on the preceding states, and differential equations describe the evolution of state. The different score components may then be considered to constitute variables in differential equations describing NK cell function and proliferation. In these equations, modelling the growth or function of NK cells, the signal from KIR-KIRL interactions will either amplify or diminish the proliferation constant depending on the input from KIR, that is the cumulative effect of activating KIR (*a*_*kir*_) or inhibitory kir (*i*_*kir*_) and missing KIR ligands (see Mathematical Supplement). Inhibitory KIR effect may be understood in terms of NK cell education. NK cells undergo education to ensure that if an individual is missing the iKIRL or has a corresponding aKIRL for their own HLA, their NK cells will not be continually activated and cause autologous tissue injury, or alternatively, if they have a high inhibitory KIR complement, they do adequately proliferate when faced with an appropriate stimulus. Education of NK cells causes them to dampen their proliferation in the former setting, and amplify it in the latter. It is logical that this education (or signal modulation in physical terms) will be proportional to the magnitude of the activating or inhibitory signals. The data presented here, support the notion that high iKIR content promotes a robust response in the face of an appropriate stimulus^31,32^ and that donor iKIR gene content may have the most significant effect on clinical outcomes.

In conclusion, KIR-KIRL interactions in an HLA-matched URD HCT setting are variable and influence both the risk of all-cause mortality and relapse risk in patients with hematological malignancies. If verified in a larger cohort of patients these findings have the potential to alter the current practice of donor selection in the HLA matched setting, and potentially in the haploidentical related donor setting as well.

## Supporting information

Mathematical Supplement

## Author Contribution Statement

EK: Designed study, collected and analyzed data, wrote the paper. RS: Analyzed data, wrote the paper. SM: collected data, wrote the paper. CC: collected data, wrote the paper. CR: collected data, wrote the paper. PK: Supervised and performed KIR typing, wrote the paper. AC: Designed study, wrote paper. AK: Wrote the paper. RR: Designed study, wrote paper. CW: Designed study, wrote the paper. RQ: Analyzed data, wrote the paper. and AT: Designed study, analyzed data, wrote the paper. AT was supported by research funding from the NIH-NCI Cancer Center Support Grant (P30-CA016059; PI: Gordon Ginder, MD).

## Conflict-of-Interest Disclosure

The authors have no relevant conflicts of interest to disclose.

## Acknowledgments

The authors gratefully acknowledge Ms. Dana Broadway for her role in managing the HLA & KIR typing of the unrelated donor transplant recipients at VCU.

