## Supplementary material for "Killer Immunoglobulin-like Receptor-Ligand Interactions Predict Clinical Outcomes Following Unrelated Donor Transplants": Mathematical Supplement

### Supplementary Materials.

#### Mathematical Supplement

There are several interesting questions raised by our model, one of which is, while weighted scores are predictive of clinical outcomes, total score in and of itself is not. It is known that NK cells express their KIR in a stochastic fashion[19], where within an individual's NK cell repertoire some NK cells express no KIR, while others express every KIR within their genotype, as well as every combination of KIR expression in-between. This implies that there may be as many as $x$ NK cell 'clones', each expressing a different KIR complement.

$$x = \left(\frac{n!}{k!(n-k)!}\right) + \left(\frac{n!}{(k-1)!(n-(k-1))!}\right) + \left(\frac{n!}{(k-2)!(n-(k-2))!}\right) + \cdots \left(\frac{n!}{(k-n)!(n-(k-n))!}\right)$$

This may be simply stated, that there are

$$x = \sum_{k}^{k-n} \left(\frac{n!}{k!(n-k)!}\right) \text{..... [7]}$$

possible NK cell clones found within every person's NK cell repertoire. In this equation, $n$ represents the number of KIR genes in the patient's genome, and $k$ represents the number of KIR genes expressed in a given NK cell clone. The sum of all possible expressed KIR gene combinations then constitutes the number of possible NK cell clones, $x$. The NK cell clone number can be calculated by taking the $n$ possible values of $k$, and calculating the number of combinations for each $k$. The model reported herein scores the KIR-KIRL interaction score on the extreme end of this spectrum, assuming that all of the known KIR are expressed on all the NK cells which may account for variable predictability within scores and score components. At this time, it is unknown what effect allogeneic transplantation will have on the distribution of KIR expression on the NK cells[28]. This is an area which needs to be prospectively explored, to determine the effect KIR expression has on KIR-KIRL interactions in the post HCT setting. This may allow more accurate weights to be ascribed to each category of the interactions in the model reported here and improve its predictive value.

The other question raised by our findings is why do inhibitory KIR have such a profound effect on clinical outcomes? To develop a quantitative understanding of KIR-KIRL interactions one may take a dynamical systems view of NK cell responses. In this system the future states of the system are dependent on the preceding states of the systems, and differential equations describe the evolution of such systems. The different score components constitute variables in these equations describing NK cell function and proliferation. One method to accomplish this includes using the logistic equation of growth which may be used to model NK cell proliferation

$$N_t = \frac{K_{NK} * N_0}{(K_{NK} - N_{t-1})(e^{-rt}) + 1}$$

In this equation: $N_0$, is the NK cell count at the outset ($N_0$=1 at $t$=1); $N_t$, is the NK cell count at time $t$ following transplant ($t$ modeled as iterations); $N_{t-1}$ represents the T cell count at the previous iteration; $K$ is the NK cell count at the asymptote (steady state conditions after infinite iterations and may be considered a proliferation constant), and represents the maximum NK cell count the system would support (carrying capacity); $r$ is the growth rate. The NK cell population, at time $t$, $N_t$, depends on the preceding T cell populations, such that

$$N_0\ [r] \rightarrow\ N_{t-1}[r] \rightarrow N_t[r] \cdots \rightarrow K_{NK}$$

In performing these calculations, $K$ is dependent on several factors. For NK cells an important variable will be the effect of KIR-KIRL, since the KIR dependent signals drives NK activity ($S_{kir}$, calculated in equation 1). The signal from the KIR (e^$S_{kir}$) will either amplify or diminish the proliferation constant depending on the input from KIR, and given the growth kinetics generally observed in immune cell proliferation, would take an exponential form. For simplicity it may be assumed that proliferation is a surrogate for activation and function.

$$N_t = \frac{(e^{S_{kir}} * K_{NK}) * N_0}{(e^{S_{kir}} K_{NK} - N_{t-1})(e^{-rt}) + 1}$$

The value $S_{kir}$ represents the cumulative effect of activating KIR ($a_{kir}$) or inhibitory kir ($i_{kir}$) and missing KIR ligands. Substituting the negative values of $i_{kir}$ and positive values of $a_{kir}$ and missing KIRL from the earlier calculations will result in shrinking or growth of the NK cell population as time passes following HCT. This becomes clear if we model the effect of the different KIR-KIRL interactions separately, in other words use the score components (activating, a; inhibitory, I; missing ligand, m) to determine signal strength, $S_{akir}$, $S_{ikir}$, and $S_{mkir}$. These equations however raise the question as to why inhibitory KIR with a negative exponent should have such a profound effect on clinical outcomes, and activating KIR with a positive exponent, such a limited impact. The answer to this lies in understanding the NK cell education and thinking of post-transplant events as a function of time following HCT.

NK cells undergo education to ensure that if an individual is missing the iKIRL or has a corresponding aKIRL for their own HLA, their NK cells will not be continually activated and cause autologous tissue injury, or alternatively, if they have a high inhibitory KIR complement, they do adequately proliferate when faced with an appropriate stimulus. Education of NK cells causes them to dampen their proliferation in the former setting, and amplify it in the latter. It is logical that this education (or signal modulation in physical terms) will be proportional to the magnitude of the activating or inhibitory signals. So while, homeostatic NK cell proliferation in response to cytokines can be modelled as a function of $K*e\hat{}S_{kir}$, a second term describing the process of NK cell education is necessary. This term makes NK cell proliferation in response to a target *inversely* proportional to the magnitude of the activating or inhibitory signals ($1/e\hat{}S_{kir}$). With inhibitory KIR this term will counteract the negative growth effect upon encountering the target and with activating KIR, dampen the signal. This relationship serves to balance the signal input with the genetically hardwired information that NK cells have to work with.

In other words, the education can render these cells inert, and drive down the whole term $K*e\hat{}S_{kir}$, if one is missing KIR Ligands for the inhibitory KIR expressed, or if the NK cells have activating KIR for KIR ligands they possess. An increase in the number of iKIR interacting with HLA molecules results in a stronger target cell response[31]. And when there are a number of inhibitory KIRs, with the NK cells getting a large inhibitory input, it may set them to a higher threshold level of activation at base line, such that when the inhibitory signal is removed the NK cell responses are augmented. This is analogous to a rubber band being imparted with potential energy as it is stretched, such that the more it is stretched, the greater the force of its rebound when it is released. Similarly, when *iKIR* with corresponding *iKIRL* are abundant, homeostatic proliferation in response to cytokines is very high, and when the inhibitory signals are turned off proliferation is rapid. So for the *iKIR* component the NK cell proliferation equation will be modified as follows

$$N_t = \frac{(1/e^{ikir}) * (e^{ikir} K_{NK}) * N_0}{((1/e^{ikir}) * e^{ikir} K_{NK}) - N_{t-1})(e^{-rt}) + 1}$$

For a missing KIR ligand component the equation will be modified

$$N_t = \frac{(1/e^{mkir}) * (e^{mkir} K_{NK}) * N_0}{((1/e^{mkir}) * e^{mkir} K_{NK}) - N_{t-1})(e^{-rt}) + 1}$$

The activating KIR will have a similar component determining equation,

$$N_t = \frac{(1/e^{akir}) * (e^{akir} K_{NK}) * N_0}{((1/e^{akir}) * e^{akir} K_{NK}) - N_{t-1})(e^{-rt}) + 1}$$

These equations together define the components of the final NK cell proliferation vector, which may be used to understand NK cell responses to stimuli (Figure X). When determining final NK cell responses, the variation in $S_{kir}$ components and the effect of education, as well as impact of cytokine mediated proliferation will have to be taken into consideration simultaneously. It is important to recognize that the NK cell proliferation in this instance is a mathematical surrogate for NK cell activation and effector functions.

Another way to understand NK cell education is if one makes an assumption that over time, NK cells have an equal basal proliferation in a cytokine-free environment. This implies that to compensate the inhibitory KIR input the proliferation coefficient will vary as a function of the KIR input (through the term *1/e^ikir*), in this instance the larger the inhibitory input with lower values of *e^Skir*, the larger the coefficient becomes with time to maintain steady state NK cell levels. Once the inhibitory influence is removed this results in exponential growth, analogous to releasing the brake on a car which has its engine revved up as it starts motion on a slope. For the activating KIR the opposite effect will hold, that education dampens the NK cell proliferation through a coefficient which lowers the basal activity. So for a high inhibitory input, $e^{-x}$, the proliferation coefficient will have to be high ($1/ e^{-x}$), and alternatively for a high activating input, $e^{x}$, the coefficient will be low ($1/e^{x}$). This also helps understand why in the model reported here, the total KIR-KIRL interaction score as calculated in Equation 1 did not predict outcomes as well as the absolute magnitude of KIR score components calculated in Equations 2-4.

This concept of KIR signal inputs applied to the NK cells to maintain steady state NK cell populations at an arbitrary basal level works well for both, when the signals are removed, so there is a commensurate rebound activation, and for the inhibitory KIR input education and for activating education. Cytokine effects are on top of this KIR mediated growth. This implies that that the proliferation coefficient is a variable quantity which starts at a certain basal value and then varies because of education, such that despite the KIR input the NK cell function may be adjusted to prevent either autologous killing or inertness in the face of a threat. In other words, to begin with proliferation coefficient is determined by the cytokine milieu and the KIR input decides NK cell function early after transplant. However, when education takes effect over time, the response to KIR input is adjusted as this process is completed. The magnitude of this effect may vary amongst the NK cell 'clones' depending on the expressed KIR complement of these cells.

**Supplementary Table 1.** Known KIR-KIR ligand interactions.

| Inhibitory | |
|---|---|
| KIR2DL1 | C2 |
| KIR2DL2 | C1 |
| KIR2DL3 | C1 |
| KIR3DL1 | Bw4 |
| KIR3DL2 | HLA A 3, A 11 |
| Activating | |
| KIR2DS1 | C2 |
| KIR2DS2 | HLA- A 11 |
| KIR2DS4 | HLA- A 11 |
| KIR2DS5 | C2 |

**Supplementary Figure 1.** NK cell alloreactivity vector and its components.

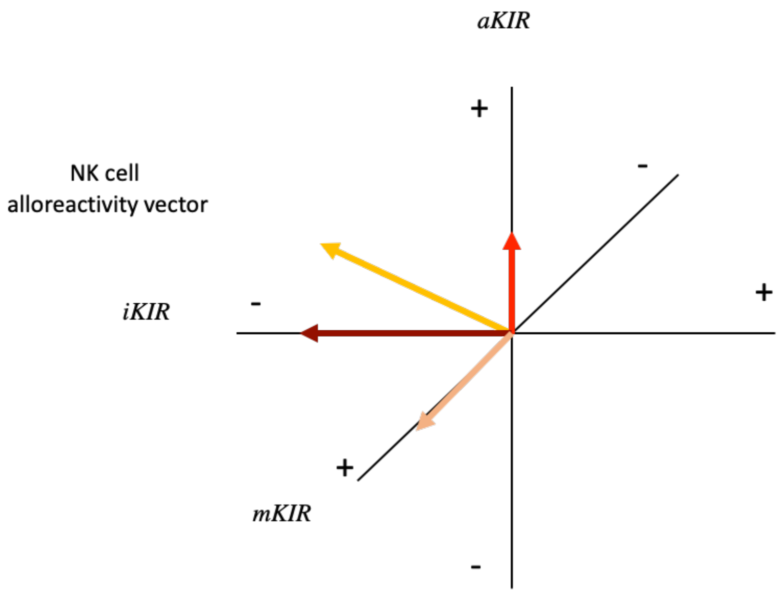
